# Concentration-dependent change in hypothalamic neuronal transcriptome by the dietary fatty acids: oleic and palmitic acids

**DOI:** 10.1101/2021.08.03.454666

**Authors:** Fabiola Pacheco Valencia, Amanda F. Marino, Christos Noutsos, Kinning Poon

**Affiliations:** Department of Biological Sciences, SUNY Old Westbury, Old Westbury NY, United States

**Keywords:** hypothalamic neurons, oleic acid, palmitic acid, proliferation, apoptosis, migration

## Abstract

Prenatal high-fat diet exposure increases hypothalamic neurogenesis events in embryos and programs offspring to be obesity-prone. The molecular mechanism involved in these dietary effects of neurogenesis are unknown. This study investigated the effects of oleic and palmitic acids, which are abundant in a high-fat diet, on the hypothalamic neuronal transcriptome and how these changes impact neurogenesis events. The results show differential effects of low and high concentrations of oleic or palmitic acid treatment on differential gene transcription. Gene ontology analysis uncovered significant gene enrichment in several cellular pathways involved in gene regulation and protein production, particularly with proliferation, migration, and cell survival. The enriched signaling pathways include Wnt, integrin, PDGF, and apoptosis, in addition endocrine function signaling pathways CCKR and GnRH. Further examination of proliferation and migration show low concentrations of oleic acid to stimulate proliferation and high concentrations of both oleic and palmitic acid to stimulate apoptosis. Oleic acid also reduced hypothalamic neuronal migration, with little effects by palmitic acid. The results show direct impact of the two most abundant fatty acids in a high fat diet to directly impact hypothalamic neuronal proliferation and migration. The results also uncovered signaling pathways affected by oleic and palmitic acid and suggest a mechanism of prenatal high-fat diet induced neurogenesis events is through these two abundant fatty acids.

## 1. Introduction

The hypothalamus is a heterogeneous brain region that regulates homeostatic processes of the body, including energy sensing in relation to hunger and satiety [1]. There are many types of neurons in the hypothalamus that contain a variation of neurotransmitters, neuropeptides, and receptors that control these homeostatic processes [2]. While the ingestion of a high-fat diet in adult animal models change the expression patterns of these neuronal regulators in the hypothalamus [3, 4], exposure to these diets during pregnancy has been widely accepted to program offspring to be more prone to ingesting these diets and becoming obese [5, 6]. Neuronal prenatal programming by a high-fat diet includes epigenetic changes [7], increased neurogenesis of orexigenic neuropeptides [6, 8], increased inflammation [9, 10], and altered neuronal patterning and connections to other brain regions [11] and to the gut [12]. The molecular mechanisms leading to these developmental changes while widely studied are still under speculation.

A large component of high-fat diets used in animal model studies is the saturated fatty acid, palmitic acid, and the monounsaturated fatty acid, oleic acid, which respectively constitutes 29% and 49% of these diets [13, 14]. These two fatty acids can promote cell proliferation and differentiation both within and outside of the central nervous system [15, 16]. The ingestion of a high-fat diet during pregnancy also increases availability of fatty acids in the placenta and to the embryo itself [17, 18]. Thus, it is possible that HFD ingestion during pregnancy increases the availability of fatty acids to the developing embryonic brain. The highly concentrated neural progenitor cell population may be disturbed by the overabundance of fatty acids and lead to changes in neurogenesis events, consequently producing hyperphagic offspring. To better examine the direct effect of fatty acids on neurodevelopmental processes, this study utilizes an immortalized rat embryonic hypothalamic neuronal cell line to examine differential gene expression changes caused by oleic and palmitic acid. The findings reveal several changes in pathways relating to the cell cycle, proliferation, apoptosis, and migration, in addition to other endocrine related processes. Further examination of hypothalamic neurons shows low concentrations of oleic acid to stimulate proliferation and high concentrations of oleic and palmitic acid to stimulate cell death while inhibiting migration. These results provide potential cellular pathways that may be involved in the effects of a high-fat diet through oleic and palmitic acids on distinct aspects of hypothalamic neurodevelopment.

## 2. Methods

### 2.1 Cell culture

Immortalized embryonic day 18 (E18) hypothalamic rat neurons (rHypoE-9) were acquired from Cedarlane (Burlington, NC). Neurons were maintained in culture using 10% FBS in 1X DMEM supplemented with 1X penicillin/streptomycin (Thermofisher Scientific, Waltham, MA). Cells were placed in a humidified 5% CO_2_ incubator at 37°C. Cells were passaged every 3- 4 days when the confluency reached ∼95%. Fatty acids (Sigma-Aldrich, St. Louis, MO) were dissolved in 100% ethanol at 1000x’s the usable concentration and diluted immediately prior to use, as previously described [14, 19, 20].

### 2.2 RNA-seq

The hypothalamic neurons were treated with 1, 10, and 100 μM oleic acid or palmitic acid for 24 hours prior to experimentation (*n* = 3), as previously described [14]. Cell culture samples were collected with RNAprotect Cell Reagent (Qiagen, Germantown, MD) and the mRNA was extracted using a Qiagen RNeasy kit (Qiagen, Germantown, MD), as previously described [21]. The yield was initially quantified with a Nanophotometer (Implen, Germany), with resulting ratios of absorbance at 260 to 280 nm of total RNA from all samples ranging between 1.90 and 2.10, indicating high purity. The mRNA samples were sent to Genewiz (South Plainfield, NJ) to determine RNA integrity (RIN > 9.9 for all samples), library preparation, quantification and quality control analysis. Genewiz also performed RNA sequencing with polyA selection using Illumina HiSeq 2×150 bp single index sequencing. Sequencing yielded libraries with an average of 34 million reads. Data was returned in FastQ formats on an external hard drive. The data analysis was performed utilizing Seawulf HPC at Stony Brook University.

### 2.3 Transcriptome Data Analysis

The RNAseq analysis was first aligned and annotated to the Rattus Norvegicus Rnor 6.0 genome using STAR v2.7.6 [22] with default settings. The output “.bam” files were then quantified using Stringtie v2.1.4 [23] to estimate transcript abundances as FPKM (Fragments Per Kilobase of exon per Million fragments mapped) which normalizes transcript expression for transcript length and the total number of sequence reads per sample. The reads were first quantified to the Rnor 6.0 reference annotation sequences, merged and requantified to the global merged transcripts. Following this, the reads were analyzed in R using DEseq2 v3.12 [24] to determine differential expression across all treatment groups. Over 90,000 genes were aligned and quantified across all experimental condition.

Gene expression analysis was performed across all concentrations of palmitic and oleic acids using Genesis [25] to select genes that were differentially expressed. The data was transformed to Log2 fold change and using 1 – Pearson correlation metric and were hierarchically clustered (HCL) for differences in expression as a function of the fatty acid or genes. To understand the biological meaning of the differentially expressed genes, the resulting gene IDs was loaded into Panther (http://www.pantherdb.org) or ENRICHR (https://maayanlab.cloud/Enrichr/) for Gene Ontology (GO) enrichment analysis. To further classify the pathways affected by each individual fatty acid treatment, manual selection of only downregulated or upregulated genes for each treatment was examined to determine enrichment across each condition. For all analyses, a threshold of *p* < 0.05 was set. Ranking of GO was computed by the combined *p*-value multiplied by the z-rank (see ENRICHR). Verification of GO analysis descriptions in addition to examining function of each individual gene was manually searched using The Rat Genome Database (https://rgd.mcw.edu/GO/). Additional function of genes was cross referenced at The Human Gene Database (https://www.genecards.org).

This analysis was repeated under only oleic acid or palmitic acid conditions to examine clustering of differentially expressed genes as a function of concentration. Lastly, the data was manually sorted, filtered using Genesis and only differentially expressed genes under all three oleic or palmitic acid conditions were selected for further GO analysis. The Volcano plots and bar graphs representing GO analysis were created using ENRICHR, and HCL plots and heatmap using Genesis.

### 2.4 Proliferation Assay

The hypothalamic neurons were seeded in a 96-well plate with 10,000 cells per well. Neurons were treated for 24 hr with 1, 10, 50, 100, 200 and 400 μM oleic acid or palmitic acid (*n* = 8 per condition). A standard curve was generated from additional neurons that were seeded at a density ranging from 50 – 50,000 cells 4 hr prior to performing the cell proliferation assay to allow for attachment. The cell proliferation assay was performed as per manufacturer’s instruction using a CyQuant kit (Thermofisher Scientific), which measures the levels of DNA via fluorophore labeling. The labeled DNA for the standards, control and treated neurons were measured at 480nm. The number of neurons were calculated based on the standard curve.

### 2.5 Cell migration scratch assay

The hypothalamic neurons were seeded in a 6-well plate with 1 x 10^6^ cells per well. Once the cells reached ∼75% confluency, a scratch was drawn across the middle of the plate with a 10 μL micropipette tip. Neurons were treated for 48 hours with 1, 10, 50, 100, and 200 μM oleic acid or palmitic acid (*n* = 8 per condition). Images were taken at timepoint 0 and again at 24 hr. The area within the scratch space was blind analyzed using ImageJ. The difference in area between 0 and 24 hr for control and treatment conditions were calculated and compared.

### 2.6 Statistical Analysis

Blind-data analysis was performed for all experiments. Statistical significance for RNA-seq data analysis was built into the programs and set at a threshold of *p* < 0.05. Statistical significance for the cell proliferation and migration assays were examined using a one-way ANOVA followed by Bonferroni *post hoc* test.

## 3. Results

### 3.1 Differential gene expression changes as a function of fatty acid treatment

Examination of 1, 10 and 100 μM oleic or palmitic acid treatment on hypothalamic neurons reveal hundreds of genes to be significantly differentially expressed in comparison to control (1 μM OA = 222, 10 μM OA = 493, 100 μM OA = 450, 1μM PA = 236, 100 μM PA = 439, 10 μM PA = 465; *p* < 0.05). Further examination of differential expression reveal overlap in many genes across 2 or more conditions (**Table 1**). To examine gene expression patterns, hierarchical clustering analysis (HCL) was performed to reveal groups of genes that show similar responses to fatty acid treatment across different conditions. The HCL analysis for all treatment groups did not show a clear clustering pattern in differential gene expression. This may be attributed to the lack of overlap in gene expression among the low to high concentrations of fatty acid in addition to the heterogenous mixture of neurons in the hypothalamus. The data was next separated into either only oleic acid condition or palmitic acid condition and reanalyzed. The resulting HCL analyses reveal two distinct clusters between 100 μM oleic and palmitic acid and 1 and 10 μM oleic and palmitic acid. (**Figure 1**). Many genes that are not differentially expressed at 100 μM concentrations are either down or upregulated at 1 and 10 μM oleic and palmitic acid and vice versa. The lack of clear patterning of differentially expressed genes across 1 and 10 μM oleic and palmitic acid indicates that each fatty acid concentration itself produces distinct effects. Gene enrichment via gene ontology (GO) analysis was performed on the 3 oleic acid or palmitic acid treatment groups. The most gene enrichment occurred under cellular processes, followed by biological process and molecular function. The top 10 processes in each of these categories show significant enrichment in pathways related to chromosomal regulation, cell cycle, and mRNA and protein processing (**FIGURE 2**). These results suggest that differences in gene expression can be attributed to a concentration effect and that fatty acids impact cellular processes involving cellular processes.

**Figure 1.**
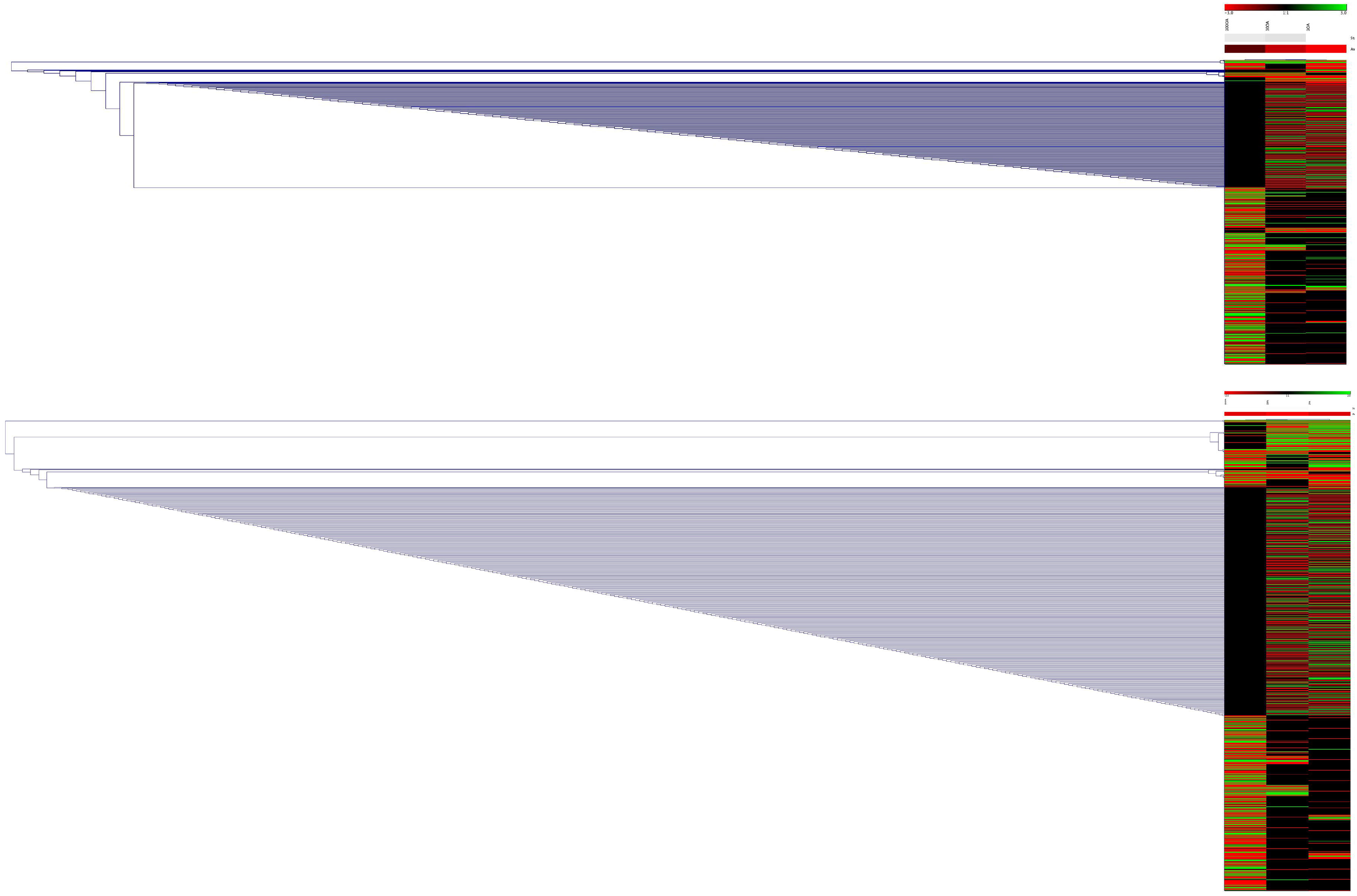
HCL analysis. Hierarchical clustering analysis of 1, 10 and 100 μM oleic acid (top) and palmitic acid (bottom) show differences between the 100 μM concentrations (left) and 1 and 10 μM concentrations (right). Many genes that are expressed in the lower concentrations are not expressed at 100 μM concentrations (black). The upregulated (green) and downregulated (red) genes do not show any further clustering.

**Figure 2.**
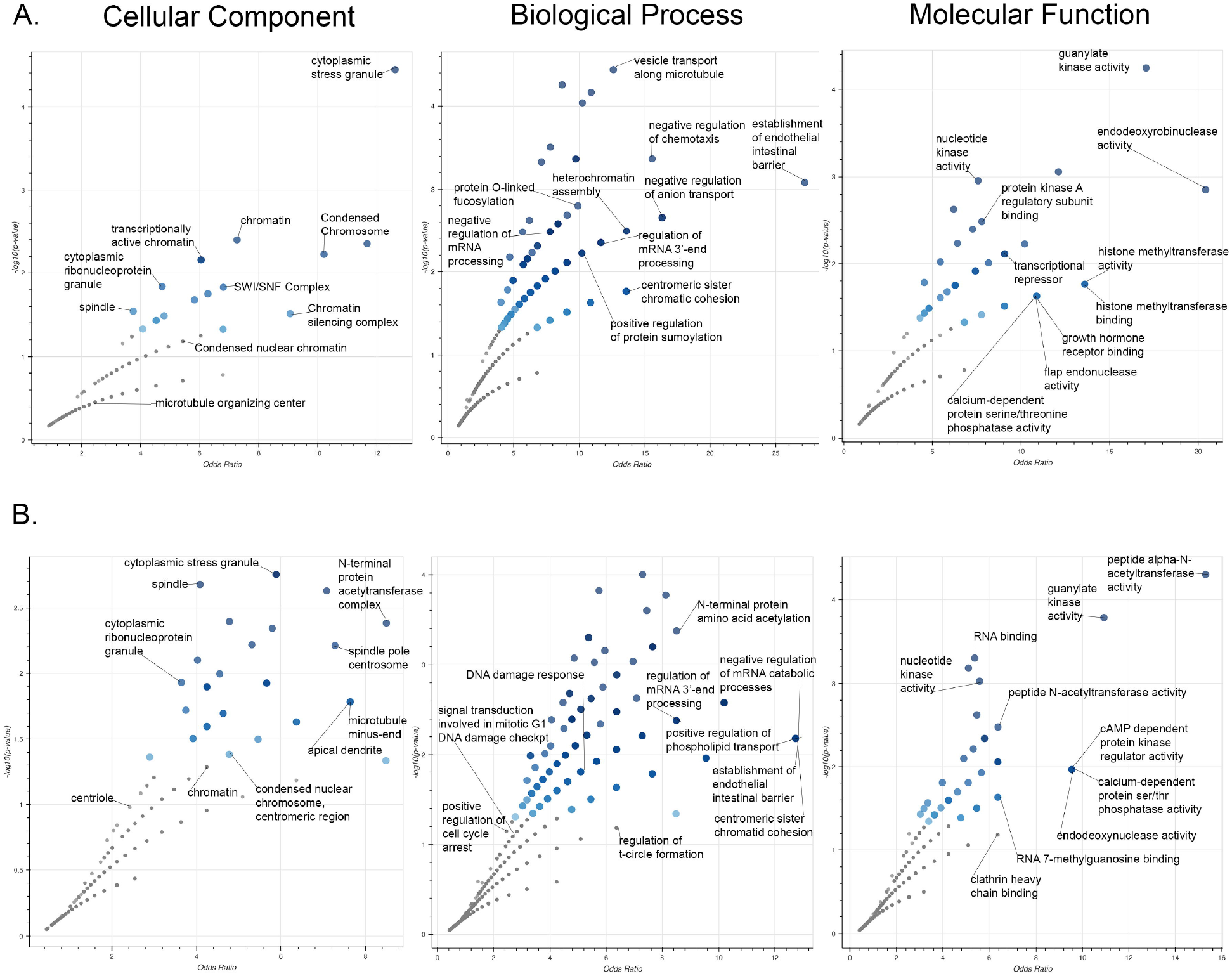
Gene ontology analysis. Gene ontology analysis of 1, 10 and 100 μM oleic acid (A) and palmitic acid (B) show significant enrichment of genes involved in cellular component, biological process, and molecular function. The top 10 significant gene sets are labeled for each process.

**Table 1.** The number of differentially expressed genes under each fatty acid condition, across two conditions, and across three conditions were determined. The boxes and numbers in red represented downregulated genes while the boxes and numbers in green represent upregulated genes.

| 3 Overlapping Positive Gene Expression |  | 13 |  |  | 11 |  |  |
| --- | --- | --- | --- | --- | --- | --- | --- |
| 2 Overlapping Gene Expression | | Oleic Acid ($\mu\text{M}$) | | | Palmitic Acid ($\mu\text{M}$) | | |
|  |  | 1 | 10 | 100 | 1 | 10 | 100 |
| Oleic Acid ($\mu\text{M}$) | 1 | 169<sup>66</sup> | 49 | 31 | 29 | 44 | 23 |
|  | 10 | 71 | 150<sup>73</sup> | 23 | 32 | 31 | 17 |
|  | 100 | 36 | 42 | 191<sup>254</sup> | 16 | 16 | 16 |
| Palmitic Acid ($\mu\text{M}$) | 1 | 49 | 59 | 31 | 172<sup>292</sup> | 32 | 35 |
|  | 10 | 50 | 71 | 34 | 45 | 290<sup>148</sup> | 17 |
|  | 100 | 37 | 92 | 30 | 37 | 41 | 321<sup>171</sup> |
| 3 Overlapping Negative Gene Expression |  | 16 |  |  | 15 |  |  |

### 3.2 Gene ontology analysis reveals trends in affected cellular pathways

The differentially expressed gene sets across varying concentrations of fatty acid suggest distinct effects on hypothalamic neurons. To examine each concentration of oleic and palmitic acid independently, GO analysis was performed for each individual treatment. The GO analysis of the upregulated and downregulated genes for each treatment revealed significant enrichment in several cellular pathways (*p* < 0.05). The top signaling pathways that were affected in at least two fatty acid conditions are: Wnt, cholecystokinin receptor (CCKR), apoptosis, platelet-derived growth factor (PDGF), integrin, and gonadotropin-releasing hormone receptor pathway (**Table 2**). The genes involved in each of the pathways were distinct across the concentrations of oleic and palmitic acid and further evidence a concentration effect on differential gene expression. The top pathways were examined in greater detail.

**Table 2.**
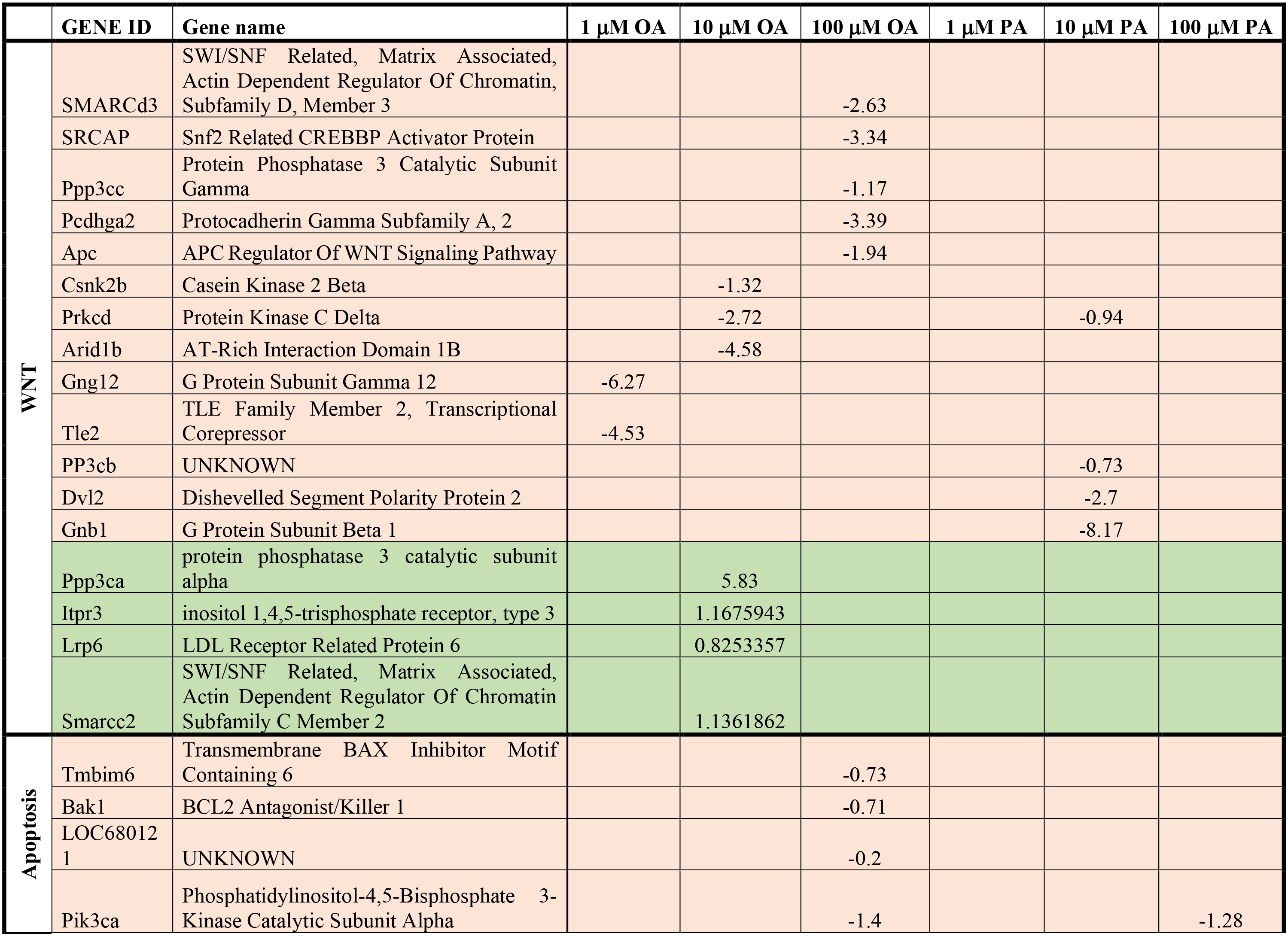
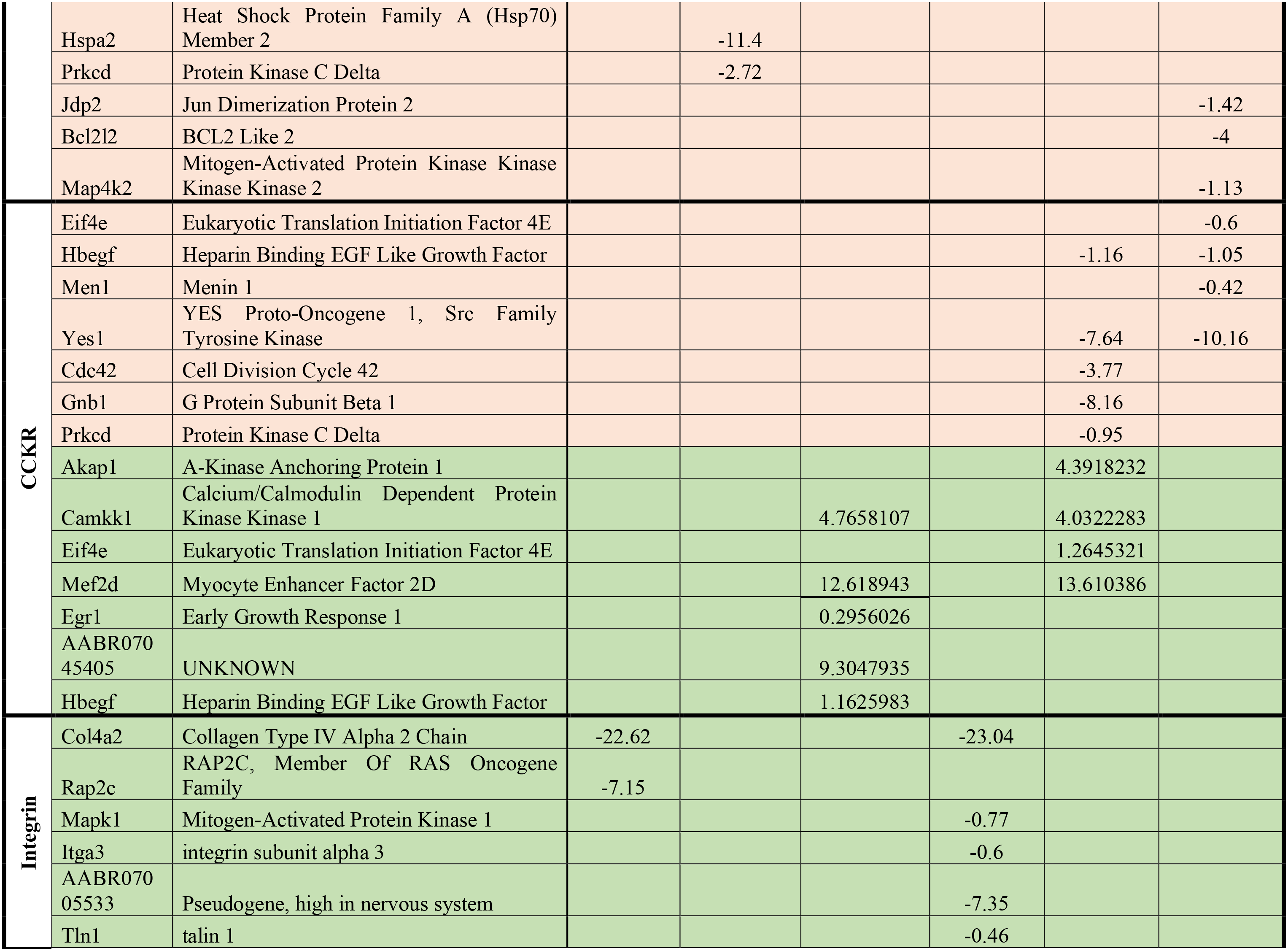
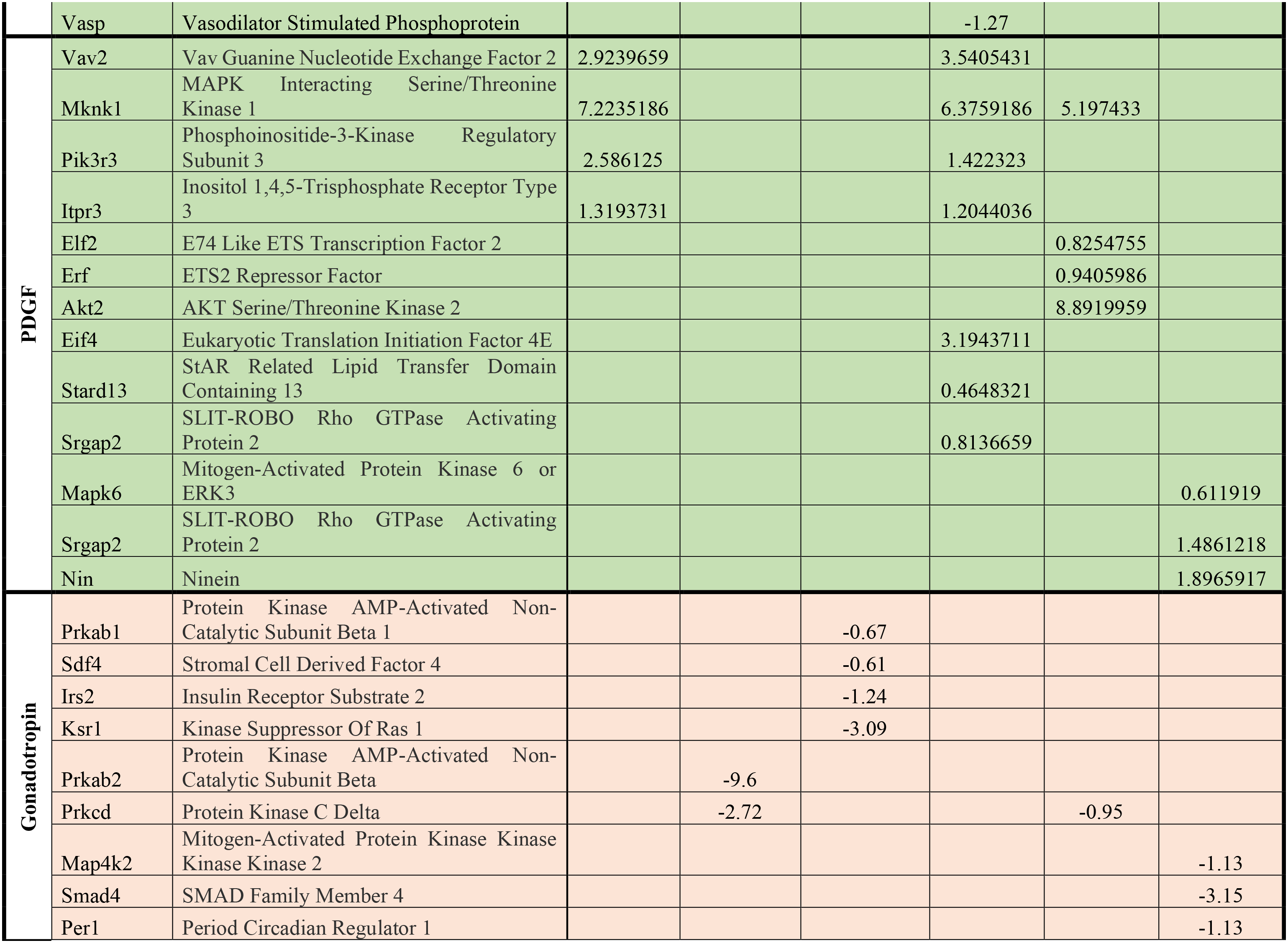
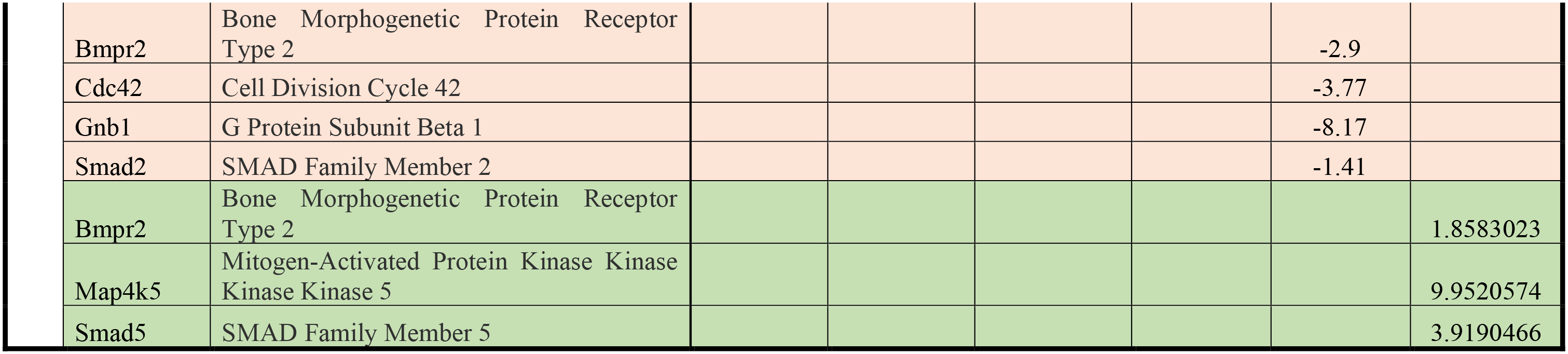
Gene Ontology (GO) analysis uncovered significant enrichment in several cellular pathways under conditions of oleic or palmitic acid (*p* < 0.05). Green are upregulated genes while orange are downregulated genes.

### 3.3 Wnt signaling pathway

The activation of canonical Wnt signaling by Wnt receptors leads to an increase in β-catenin signaling and changes gene transcription, resulting in cell proliferation and cell survival [26, 27]. Some Wnt receptors include the G-protein coupled receptor *FRIZZLED* and *LRP5/6* [28], in addition to a few tyrosine receptor kinases (*RYR* and *ROR*; **FIGURE 3**) [29, 30]. Several g-protein subunits and kinases were downregulated at 1 or 10 μM oleic acid or palmitic acid treatment (*Csnk2b, Prkcd, Gng12, Gnb1*) or upregulated at 10 μM oleic acid (*Lrp6*) and at 100 μM palmitic acid (*Csnk2a1*). Genes associated with regulating Wnt signaling were also changed. Genes that inhibit Wnt signaling [31, 32] were found to be upregulated at 100 μM palmitic acid (*Smad5, apc, Tle4*) and downregulated at 1 and 10 μM oleic acid (*apc, tle2, and dvl2*). Other changes involve the SWI/SNF (switch/sucrose nonfermentable) family of protein complexes, which activates Wnt signaling [33]. The SWI/SNF family of proteins acts as a switch in changing chromatin structure and forms large complexes to regulate gene transcription [34]. This complex is required for stem cell proliferation [35], including neuronal development [36, 37]. The SWI/SNF family of genes were downregulated at 10 and 100 μM oleic acid (*Smarcd3, SRCAP, Ppp3c, Pcdhga2, Arid1b*) while *Smarcc2* was the only upregulated gene at 10 μM oleic acid. Interestingly, genes that are involved in Ca^2+^ signaling were increased with 10 μM oleic acid (*PPP3ca, Ppp3cc, Itpr3*) and may play a role in the activation of the noncanonical Wnt activated gene transcription pathway [38]. Two genes that were upregulated at 100 μM palmitic acid had unknown function (*AC109100, Celsr2*). These gene expression changes show activation of Wnt signaling at low oleic acid concentrations suggesting cell proliferation and survival and the opposite effect at high oleic and palmitic acid concentrations.

**Figure 3.**
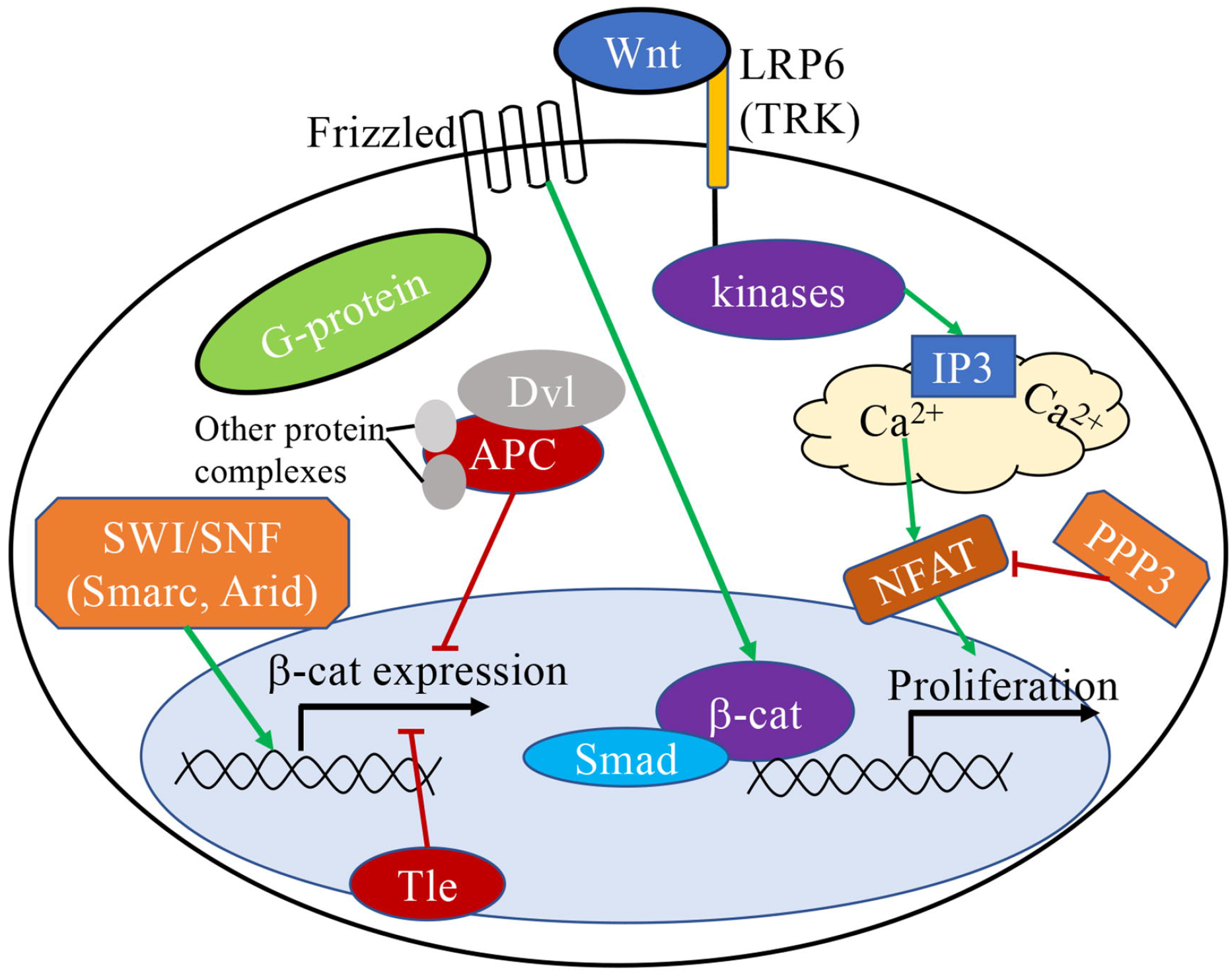
A potential mechanism of oleic and palmitic acid effects on the genes involved in the Wnt signaling pathway.

### 3.4 CCKR signaling pathway

Cholecystokinin (CCK) is a peptide released from the small intestine that signals satiety and senses lipids in the gut [39]. In the hypothalamus, neurons from the nucleus of solitary tract also contain CCK neurons that project into the hypothalamus to control hunger and satiety signaling [40]. Thus, it is not surprising that this pathway arose in GO analysis of fatty acid treatment on hypothalamic neurons. The CCK signaling pathway also overlaps with apoptosis and Wnt signaling pathways [41], and suggests that the overlap in hunger and satiety signaling with cell proliferation and apoptotic pathways may be a side effect of metabolic signaling. The pathways leading to CCK and CCKR transcription and the downstream pathways of CCKR signaling are complex (**FIGURE 4**). At 10 and 100 μM palmitic acid, several genes involved in pathways leading to gene transcriptional changes were downregulated (10: *Hbegf, Yes1, Cdc42, Gnb1, Orkcd*; 100: *Hbegf, Eif4e, Men1, Yes1*). At 10 μM palmitic acid (*Akap1, Camkk1, Eif4e, Mef2d*) and 100 μM oleic acid (*Camkk1, Egr1, Hbegf, Mef2d*), several genes involved in calcium signaling pathways were upregulated. G protein beta subunit 1 (*Gnb1*) is highly downregulated at 10 μM palmitic acid and may be coupled to CCKR receptors to activate downstream signaling. Knockdown of this G-protein subunit has been implicated in the development of obesity [42]. CCK can directly activate the eukaryotic initiation transcription factor 4 (*Eif4e*) to initiate translation [43] and is upregulated at 10 μM but down regulated at 100 μM palmitic acid. Protein kinase C delta (*Prkcd*) and YES proto-oncogene 1 (*YES1*) are downstream of both CCKR and EGFR signaling and are downregulated with 10 μM palmitic acid treatment. This pathway may be involved in the transcription of *Hbegf* [44]. Other transcription factors that were upregulated at 10 μM palmitic acid and 100 μM oleic acid include myocyte enhancer factor 2D (*Mef2d*), A-kinase anchoring protein 1 (*Akap1*), and early growth response 1 (*Egr1*). These proteins are involved in cell survival, proliferation, and migration processes [45–47]. One gene that was upregulated has not been previously identified (*AABR07045405*). These results suggest that at high concentrations of fatty acids, cellular pathways leading to cell survival are inhibited while at 10 μM oleic acid and palmitic acid there is both up and downregulation of genes involved in proliferation and migration.

**Figure 4.**
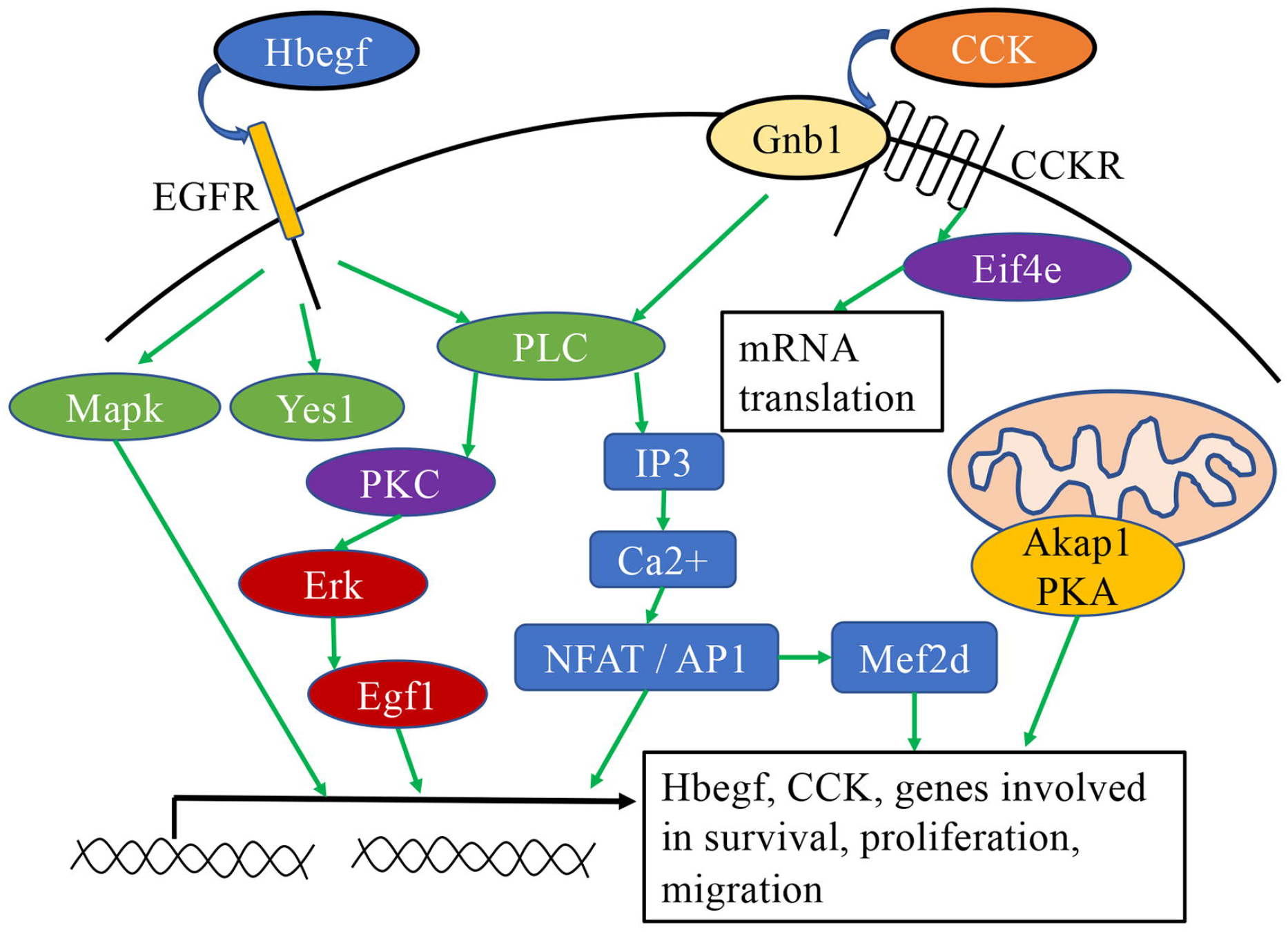
A potential mechanism of oleic and palmitic acid effects on the genes involved in CCKR signaling pathway.

### 3.5 Apoptosis signaling pathway

The classical apoptosis pathway results in the activation of caspase cascades that leads to irreversible apoptosis [48]. A method of activation of apoptosis is through the mitochondrial *Bax/Bak* receptors (**Figure 5**). Activation *Bax/Bak* stimulated by cellular damage or stress results in the release of cytochrome c, which initiates the caspase pathway [49]. The *Bcl2* and *TMBim6* family of proteins under normal conditions inhibits the activation of *Bax/Bak*. At 100 μM palmitic acid, *Bcl2l2* (Bcl-w) is highly downregulated, suggesting increased activation of the apoptotic pathway. Upstream transcriptional activation of *Bcl2l2* expression include β-catenin [50], which is also downregulated at 100 μM palmitic acid and suggests overlap with Wnt signaling. *Pik3ca* is also downregulated at 100 μM palmitic acid, with downstream activation of *Akt* signaling to also activate *β-catenin* driven *Bcl2* gene expression [51, 52]. Downregulation of *Map4k2* at 100 μM palmitic acid can inhibit *Bcl2l2* through *JNK* signaling [53]. Jun dimerization protein 2 (*Jdp2*) is a transcription factor involved in cell survival [54] and was also downregulated at 100 μM palmitic acid.

**Figure 5.**
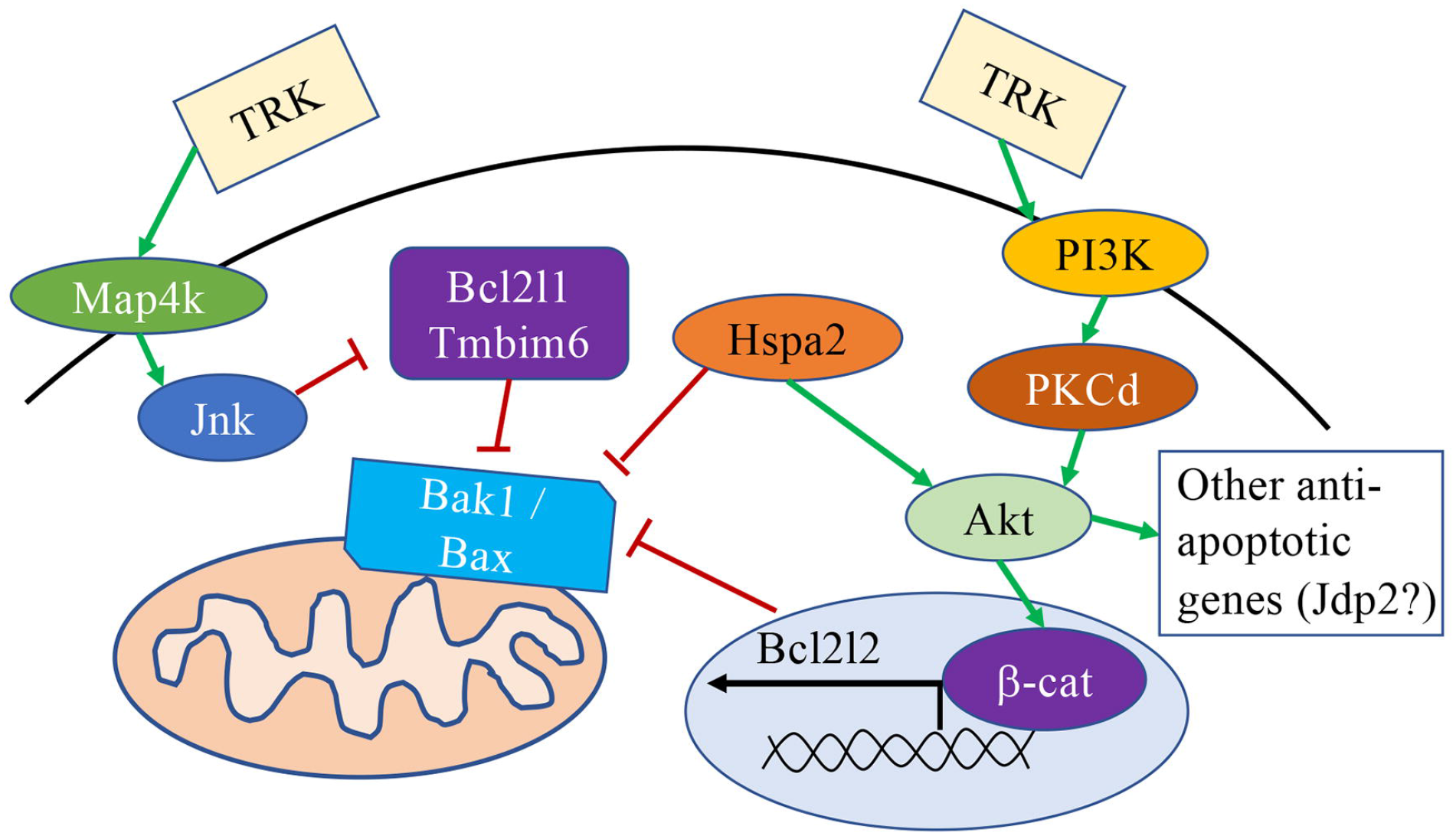
A potential mechanism of oleic and palmitic acid effects on the genes involved in the apoptosis signaling pathway.

Similar gene expression changes occurred at 100 μM oleic acid, with downregulation of *Tmbim6, Bak1* and *pik3ca*. *Prkcd* was also downregulated at 10 μM oleic acid and is downstream of pik3ca anti-apoptotic cell signaling [55]. Heat shock proteins have also been implicated in anti-apoptotic pathways, including activation of *Akt* and inhibition of *Bax/Bak* [56, 57], and one isoform (*Hspa2*) was greatly downregulated at 10 μM oleic acid. The downregulation of genes involved in anti-apoptotic pathways and upregulation of apoptosis signaling at 100 μM oleic and palmitic acid concentrations show high levels of fatty acids to be toxic to hypothalamic neurons.

### 3.6 Integrin signaling pathway

Integrins are a group of receptor proteins that interact with the extracellular matrix to induce signaling cascades involved in cell proliferation and migration processes [58]. Integrin receptor signaling can activate MAP kinase cascades and gene expression changes [59] of proteins that regulate the cell cycle (**FIGURE 6**) [58]. Integrin receptors can also be activated through an inside-out reverse signaling mechanism [59, 60]. One pathway of inside-out activation is through rap gtpases (*Rap2c*), which recruits talin (*tln1*) to *PIP3*, resulting in changes in intracellular Ca2+ levels [61]. Vasodilator stimulated phosphoprotein (*vasp*) similarly induces inside out activation of integrin receptors [62]. Extracellular matrix proteins such as collagen type IV α2 chain (*Col4a2*) can bind to integrin receptors to initiate gene transcription of *PI3K-Akt* and promote cell proliferation in tumors [46]. Several genes in the integrin signaling pathway were downregulated at both 1 μM oleic acid (*Col4a2, Rap2c*) and palmitic acid (*Cal4a2, Mapk1, Itga3, Tln1, Vasp*) treatment. The downregulation of integrin signaling pathway proteins suggest a decrease in proliferation [63, 64] and changes in migration. However, integrin signaling is complex and thus migratory processes cannot be inferred from the differentially expressed genes alone [65].

**Figure 6.**
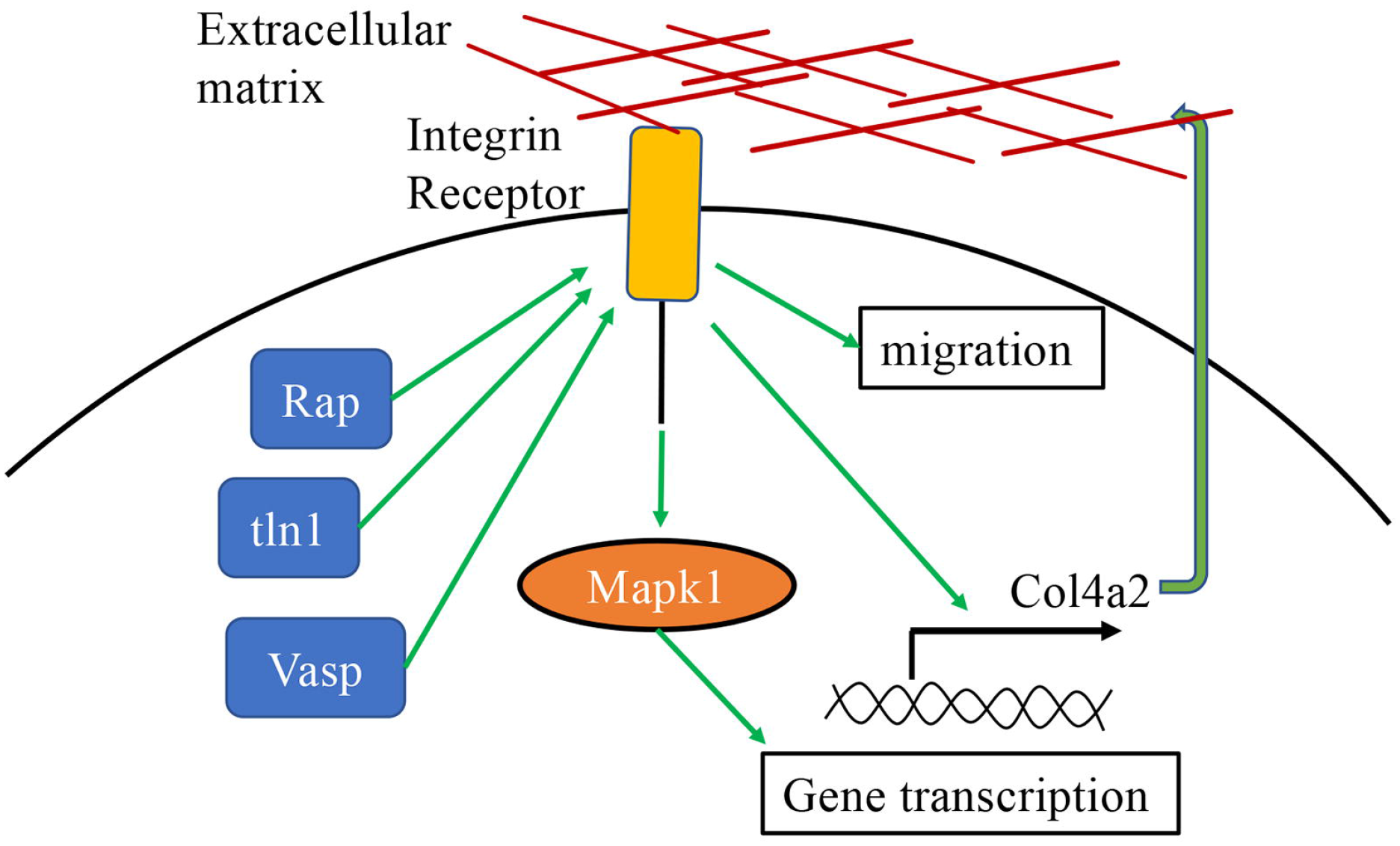
A potential mechanism of oleic and palmitic acid effects on the genes involved in the PDGF signaling pathway.

### 3.7 PDGF signaling pathway

Platelet-Derived Growth Factor (PDGF) signaling pathway regulates a diverse set of functions, including proliferation, cell survival, differentiation, modulation of receptors, and the development of specific neuronal subtypes (**FIGURE 7**) [66]. The activation of tyrosine receptor kinases is important for neurogenesis processes in the brain [66]. At all three concentrations of palmitic acid and at 1 μM oleic acid, several genes involved in cell migration processes were upregulated. The vav guanine nucleotide exchange factor 2 (*Vav2*) found to be upregulated at 1 μM oleic acid and 1 μM palmitic acid is downstream of PDGF signaling and targets *RhoA*, *CDC42*, and *RAC* to change gene transcription [67]. MAPK interacting serine/threonine kinase 1 (*Mknk1*) is upregulated in all but 100 μM palmitic acid and activates *Eif4* mediated mRNA translation of proteins involved in migration [68, 69]. Other genes downstream from PDGF activation that were upregulated at 1 μM oleic acid are phosphoinositide-3-kinase regulatory subunit 3 (*Pik3r3 or PIP3*) and inositol 1,4,5-trisphosphate receptor type 3 (*Itpr3*). Downstream *Pik3r3* is AKT serine/threonine kinase 2 (*Akt2*) and E74 like ETS transcription factor 2 (*Elf2*), which are upregulated at 10 μM palmitic acid. Other genes downstream from PDGF activation that were upregulated at 1 μM palmitic acid include StAR related lipid transfer domain containing 13 (*Stard13*), which is downstream from β-catenin signaling, and *Srgap2* that is involved in stimulating cell migration [70]. At 100 μM palmitic acid, *Mapk6, Srgap2*, and *Nin* are upregulated. *Mapk6* is involved in neuronal morphogenesis and spine formation while negatively regulating cell proliferation [71, 72]. *Nin* is involved in microtubule formation during migration and in stimulating axogenesis [73]. These selective pathways suggest that palmitic and oleic acids play a role in migration.

**Figure 7.**
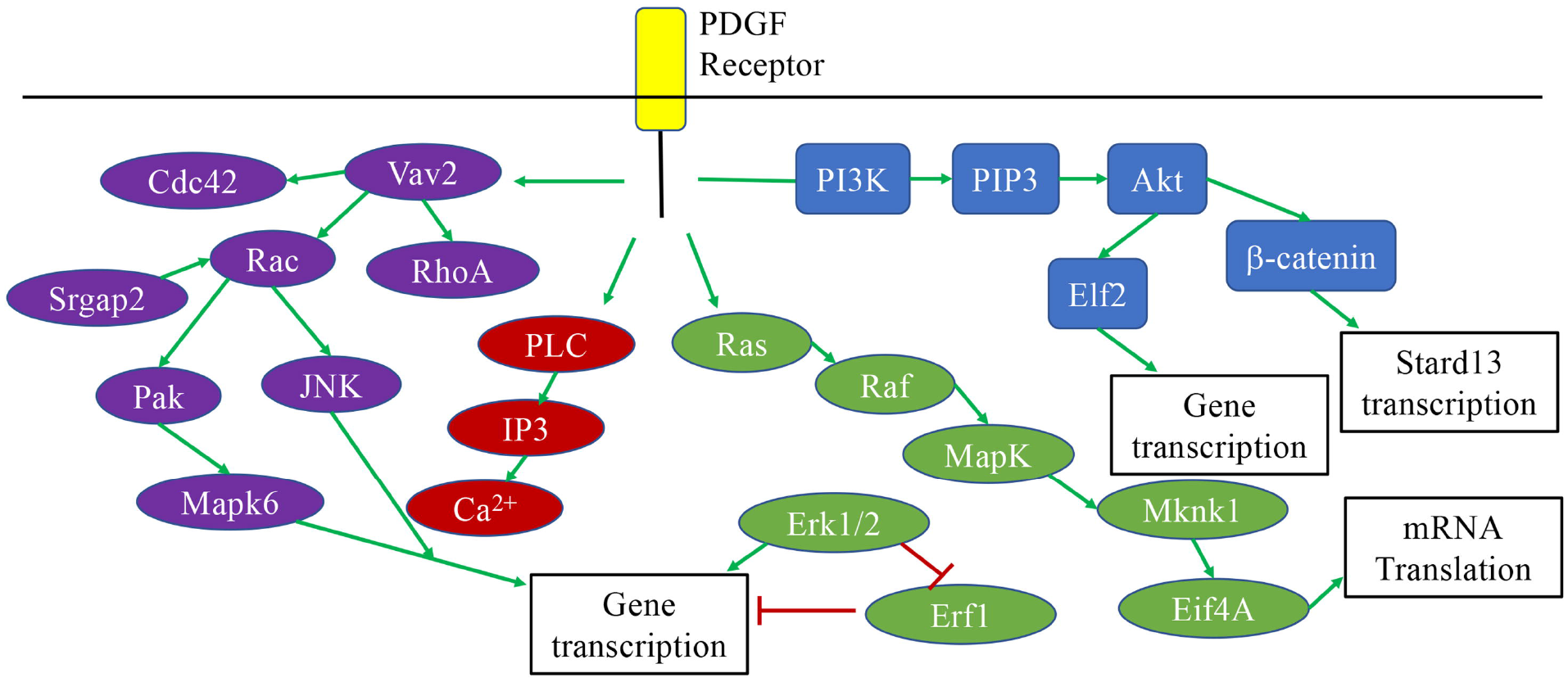
A potential mechanism of oleic and palmitic acid effects on the genes involved in the integrin signaling pathway.

### 3.8 Gonadotropin-releasing hormone signaling pathway

Gonadotropin-releasing hormone (GnRH) from the hypothalamus stimulates the release of follicle stimulating hormone and luteinizing hormone from the pituitary [74]. Several neuropeptides that regulate food intake either co-express with or are connected to GnRH neurons [75–78]. At 10 and 100 μM oleic and palmitic acid, several genes involved in gonadotropin signaling or gonadotropin gene expression were downregulated (*Prkab1, Prkab2, Prkc, Sdf4, Irs2, Ksr1, Map4k2, Smad2, Smad4, Per1, Bmpr2, Cdc42, Gnb1*). Only three genes were upregulated at 100 μM palmitic acid (*Bmpr2, Map4k5, Smad5*). The protein kinase AMP-activated non-catalytic subunit beta 1 (*Ampk*) and insulin receptor substrate 2 (*Irs2*) are expressed in hypothalamus and acts as nutrient sensors through control of orexigenic neuropeptides [76, 78, 79] and as stimulants of gonadotropin release [80, 81]. The SMAD family, period circadian regulator 1 (*per1*), *Cdc42*, and *Mapks* are downstream from GnRH signaling [82, 83]. *Smads* and bone morphogenetic protein receptor 2 (*Bmpr2*) are also involved in the transcriptional activation of GnRH [84]. These results are consistent with fatty acids modulating the activity of hypothalamic neurons involved in controlling ingestive behavior in addition to sex hormone production.

### 3.9 Differentially expressed gene overlap in oleic and palmitic acid treatment

The individual GO analysis revealed overlap in cellular pathways affected by the individual fatty acids. Using Panther, only differentially expressed genes across all three oleic acid or palmitic acid concentrations were examined and 27 genes returned matches for oleic acid and 36 genes returned matches for palmitic acid. The HCL analysis revealed 4 clusters with oleic acid treatment and 3 clusters of genes with palmitic acid treatment (**FIGURE 8**). The limited number of genes did not yield significant enrichment in any pathway during GO analysis but may be examined in future studies on modulation of neuronal function.

**Figure 8.**
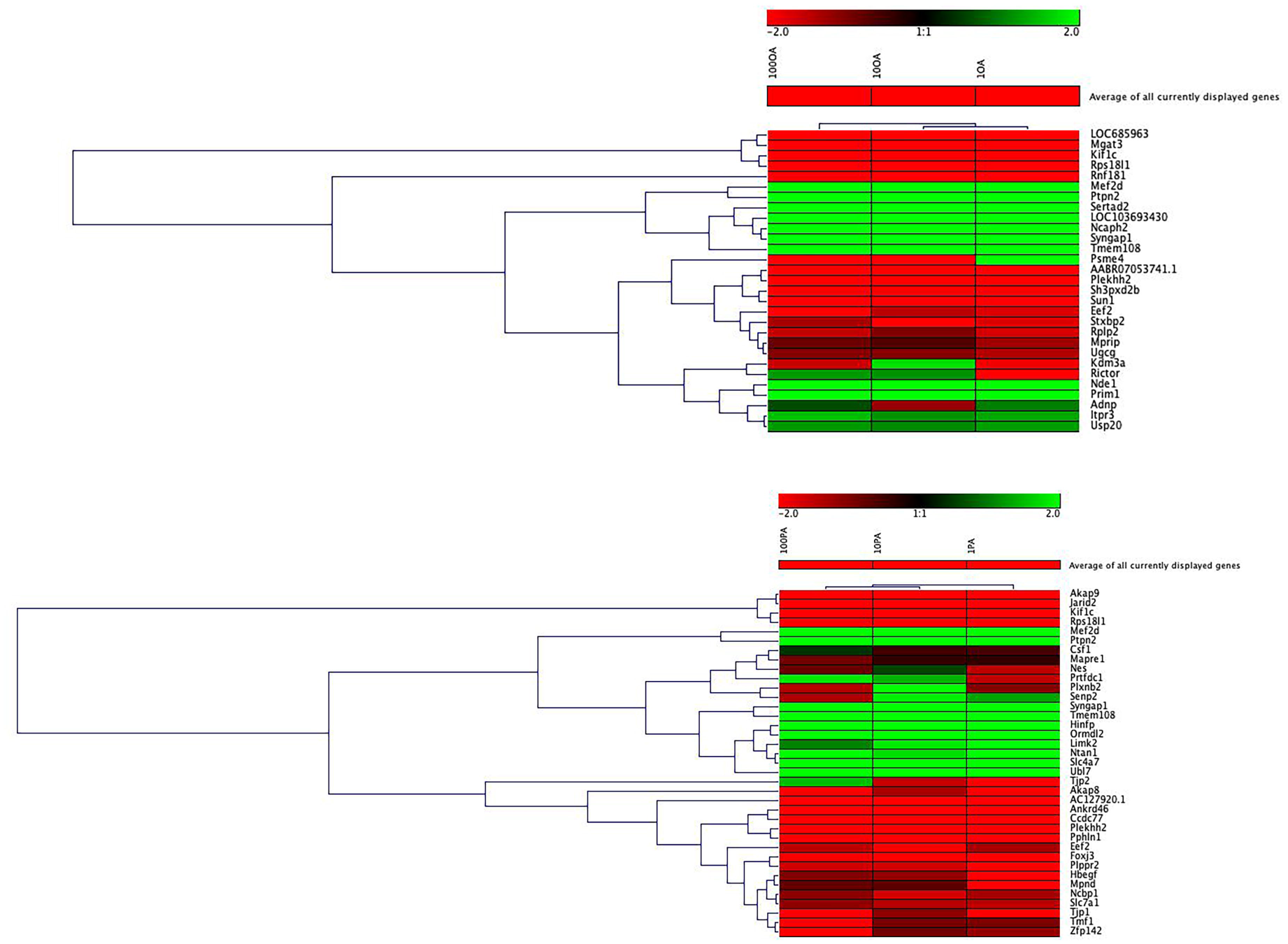
HCL analysis. Hierarchical clustering analysis of genes that are expressed under all three conditions of oleic acid treatment (top) or palmitic acid treatment (bottom). Four clusters are present with oleic acid and three with palmitic acid treatment.

### 3.10 Oleic and palmitic acid stimulates proliferation and cell death and inhibits migration

Based on the transcriptome analysis, both oleic and palmitic acid is involved in the progression of cell cycle processes, migration and apoptosis. To examine whether these fatty acids affect hypothalamic neuronal proliferation and death, a cell proliferation and viability test was performed. Treatment with 1, 10, 50, 100, 200 and 400 μM oleic or palmitic acid for 24 hr had a significant effect on the number of hypothalamic neurons (*F* (12, 91) = 65.49, *p* < 0.01). Bonferroni *post hoc* tests reveals significant change in the number of neurons at all concentrations of oleic acid and at 1, 50, 200 and 400 μM palmitic acid (**Table 3; Figure 9B**). While both low concentrations of oleic (1, 10 μM) and palmitic acid (1 μM) stimulated cell proliferation, concentrations greater than 50μM reduced cell viability. Visual examination showed that many cells died with 400 μM oleic and palmitic acid treatment.

**Figure 9.**
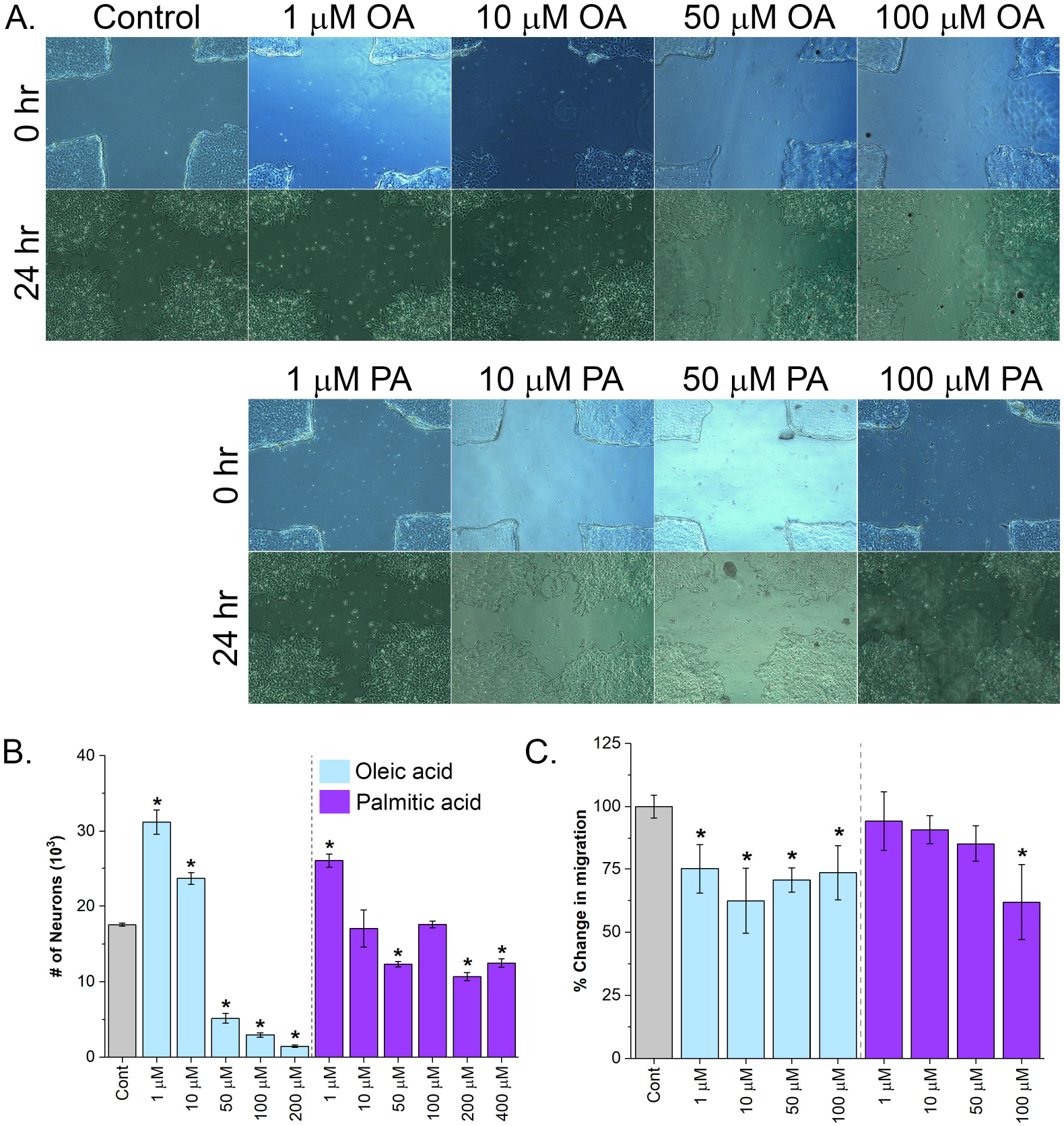
Proliferation and migration. Proliferation and migration analysis of hypothalamic neurons reveals significant effects by oleic and palmitic acid. A. Representative images of the scratch assay used to measure migration of the hypothalamic neurons over 24 hr. B. The 1 and 10 μM oleic acid significantly increased proliferation. 50 μM and higher concentrations of oleic acid induced significant cell death. Similar reduction in neuronal number was found at 50, 200 and 400 μM palmitic acid. C. Treatment with all concentrations of oleic acid and only at 100 μM palmitic acid significantly reduced migration of neurons. \**p* < 0.05.

**Table 3.** Bonferroni *post hoc* test for the proliferation assay reveal statistical significance on proliferation across all concentrations of oleic acid (OA) and palmitic acid (PA) treatment except at 10 and 100 μM palmitic acid.

|  | Proliferation |  | Migration |  |
| --- | --- | --- | --- | --- |
| **Treatment** | ***t*(14) = value** | ***p*-value** | ***t*(14) = value** | ***p*-value** |
| 1 mM OA | 7.96 | < 0.01 | 3.67 | < 0.05 |
| 10 mM OA | 3.57 | < 0.05 | 5.55 | < 0.01 |
| 50 mM OA | 7.22 | < 0.01 | 4.34 | < 0.01 |
| 100 mM OA | 8.50 | < 0.01 | 3.9 | < 0.01 |
| 200 mM OA | 9.38 | < 0.01 | - | - |
| 400 mM OA | 10.64 | < 0.01 | - | - |
| 1 mM PA | 4.94 | < 0.01 | 0.84 | 1 |
| 10 mM PA | 0.30 | 1 | 1.35 | 1 |
| 50 mM PA | 3.04 | < 0.05 | 2.18 | 1 |
| 100 mM PA | 0 | 1 | 5.63 | < 0.01 |
| 200 mM PA | 4.01 | < 0.01 | - | - |
| 400 mM PA | 2.96 | < 0.05 | - | - |

The fatty acids were also examined for its ability to affect hypothalamic neuronal migration using a simple scratch assay. Treatment with 1, 10, 50, 100 and 200 μM oleic or palmitic acid for 24 hr reveal significant effects on migration (*F* (8, 63) = 8.37, p < 0.05). (**Figure 9A, C**). Bonferroni *post hoc* test reveals a significant decrease in migration at all concentrations of oleic acid and a significant decrease at 100 μM palmitic acid (**Table 3**). The measured decrease in migration at the high concentrations of fatty acid may be due to the increase in cell death. These results suggest that oleic acid inhibits while palmitic acid does not impact hypothalamic neuronal migration.

## 4. Discussion

The molecular process governing maternal high-fat diet-induced neurogenesis events in the brain have been a focus on understanding prenatal programming of ingestive behavior. In these offspring, the hypothalamus has an increased number of orexigenic peptide neurons that induces a hyperphagic phenotype [6, 8]. A potential mechanism involved in high-fat diet outcomes on hypothalamic neurogenesis events directly involve the fatty acids, oleic and palmitic acids.

### 4.1 Distinct concentration-dependent effects of oleic and palmitic acid on differential gene expression

The initial analysis of the combined fatty acid groups displayed nonspecific gene clusters that could be attributed to both a concentration effect and differences in fatty acid pathway activation. Examining only oleic or palmitic acid alone revealed clustering differences between 100 μM to 1 and 10 μM, without any further emerging patterns. The individual GO pathway analysis revealed each fatty acid to change the expression of genes involved in specific aspects of each signaling cascade while maintaining the findings of the HCL analysis. For each concentration of fatty acid, only a particular subset of genes involved in each pathway was found to be differentially expressed. Low concentrations of oleic or palmitic acid stimulated anti-apoptotic targets and inhibited apoptotic targets in Wnt and apoptosis signaling pathways while the opposite effect was observed at primarily 100 μM concentrations. A single fatty acid can have a concentration effect to impact different aspects of a signaling cascade and would account for the diverse programming effects observed during the neurodevelopmental period.

### 4.2 High fat diet-induced neurogenesis is partially mediated by fatty acids

Neurogenesis from stem cell populations involves proliferation, differentiation and migration steps for proper formation of the embryonic brain [85]. This delicate process can be disturbed and impact the modeling of brain tissue, particularly during exposure to excessive prenatal dietary fat [6]. The RNAseq analyses reveal enrichment in several pathways involved in proliferation, migration, and apoptosis signaling. The results from this study show oleic acid to have a concentration dependent increase in hypothalamic neuronal proliferation followed by cell death. These results are in line with studies showing prenatal oleic acid exposure to increase overall brain neurogenesis [86] while high concentrations stimulate cell death [87]. Palmitic acid has also previously been shown to induce cell death in PC12 cells [20] and is similar to the findings in this study. In hippocampal neural progenitor cells, there is an increase in the production of fatty acids and a decrease in fatty acid oxidation during proliferation while the reverse is true during cell quiescence [16, 88]. This suggests that changes in fatty acid metabolism and levels impacts neuronal proliferation processes and lead to changes in the phenotypic development of neurons.

There is limited evidence that supports or refutes oleic and palmitic acid’s role on neuronal migration. Both fatty acids can either stimulate or inhibit migration and may be dependent on the concentration and the cellular subtype examined [89–91]. The results from this study show oleic acid to reduce neuronal migration with no effect by palmitic acid. These results suggest the importance of oleic acid in promoting hypothalamic neuronal proliferation but not on migration. Palmitic acid has previously been shown to be cytotoxic to neuronal stem cells while low concentrations can stimulate differentiation (Wang Z, Hao A, 2014). The effect of palmitic acid on hypothalamic neurogenesis processes is not as clear but may involve differentiation. This last aspect of neurogenesis could not be examined in this study because the hypothalamic neurons have already differentiated.

### 4.3 Evidence for oleic and palmitic acid involvement in orexigenic neuropeptide signaling pathways

The RNA-seq did not detect any changes to the typical fat-sensing hypothalamic orexigenic neuropeptides, such as enkephalin and galanin. The hypothalamus typically begins to express these neuropeptides at embryonic day 17-18 and may be too early to detect the fat-sensing neuropeptides [21]. However, the enrichment of genes involved in Wnt signaling pathways may be upstream effectors of neuropeptide expression. The three concentrations of oleic acid treatment had impacted gene expression changes in proteins downstream of Wnt signaling, SWI/SNF and Arid1 (**Table 2; Figure 3**). The inactivation of SWI/SNF and Arid1 leads to the activation of the YAP/TAZ/TEAD transcriptional regulator complex to increase proliferation [92]. Meanwhile, prenatal high-fat diet exposure decreases the activity of the transcriptional regulators YAP and TEAD, with this decrease to stimulate the expression of the orexigenic neuropeptide, enkephalin [93]. This evidence suggests high-fat diet induced neurogenesis events may be mediated by fatty-acid effects on the Wnt signaling pathway, presumably to increase proliferation of hypothalamic orexigenic peptide neuronal precursors and to later increase the neuropeptide levels.

Treatment of 10 and 100 μM oleic and palmitic acids also impacted the GnRH signaling pathway. The hypothalamic GnRH neurons coexpress or contact with orexigenic peptide neurons, such as neuropeptide Y and galanin, [75–78]. These hypothalamic hormone pathways regulate endocrine function and are impacted by dietary changes. There is abundant evidence that obesity negatively impacts successful fertility outcomes and prenatal obesity and high-fat diet exposure to influence offspring pubertal development and fertility [94–96]. The results of this study also suggest endocrine programming is in part mediated by high fatty acid levels that are increased during obesity and excessive dietary fat intake.

While activation of CCKR signaling pathway leads to changes in gene transcription involving cell survival, proliferation and migration, CCKR activation is also heavily linked to satiety and decreased eating [97]. Genetic knockout of CCKR1 in rats have an obesogenic and hyperphagic phenotype [98]. There is also evidence linking direct activation of CCKRs to stimulate orexigenic peptide neurons in addition to coexpression with feeding neuropeptides [99, 100]. During the fed-state, CCK is released by the gut to signal satiety in the hypothalamus [99, 100]. The differential expression of genes involved in the CCKR signaling pathway suggests high concentrations of fatty acids lead to dysfunctional gut-brain signaling in the intact organism.

### 4.4 Oleic and palmitic acids mediate high-fat diet changes in signaling pathways associated with metabolism

Other signaling pathways that were uncovered by oleic and palmitic acid treatment on hypothalamic neurons overlap with outcomes from *in vivo* high-fat diet studies. The Wnt signaling pathway has been previously shown to be involved in high-fat diet effects on several physiological systems, including bone, adipose tissue, vascular tissue, and brain (Bagchi DP, Macdougald OA, 2020; Chen N, Wang J, 2018). Changes or mutations with the Wnt signaling cascade leads to metabolic disorders, including targets uncovered in this study (Chen N, Wang J, 2018). Integrin signaling in adipose tissue has also been associated with metabolic effects resulting from high-fat diet intake, particularly in the development of obesity-induced diabetes (Williams AS, Wasserman DH, 2016; Ruiz-Ojeda FJ, Ussar S, 2021). Excessive ingestion of high-fat diets has been linked to increased activation of apoptosis signaling pathways in the hypothalamus (Moraes JC, Velloso LA, 2009). Excessive high-fat diet ingestion and metabolic changes are also mediated by receptor tyrosine kinases, with changes in PDGF signaling pathway as one example (Zhao M, Svennson KJ, 2020). These results highly suggest high-fat diet effects are primarily mediated by oleic and palmitic acids.

### 4.5 Conclusions

The results from this study reveal several potential methods of oleic and palmitic acid effects on neurogenesis events, particularly involving signaling pathways that intersect with orexigenic neuropeptides and cell survival. Further studies examining each distinct pathway on neuropeptides may further elucidate the molecular mechanisms of prenatal high-fat diet exposure on neurogenesis events.

## Acknowledgements

This work was funded by a SUNY Old Westbury Faculty Development Grant and from the institute for cancer research and education (ICaRE) at SUNY Old Westbury. The authors would like to thank Stony Brook Research Computing and Cyberinfrastructure, and the Institute for Advanced Computational Science at Stony Brook University for access to the high-performance SeaWulf computing system, which was made possible by a $1.4M National Science Foundation grant (#1531492).

## Notes

### Competing Interest Statement

The authors have declared no competing interest.

